# Chromatin features constrain structural variation across evolutionary timescales

**DOI:** 10.1101/285205

**Authors:** Geoff Fudenberg, Katherine S. Pollard

**Affiliations:** Gladstone Institute of Data Science and Biotechnology, San Francisco, California, USA.; Department of Epidemiology & Biostatistics, Institute for Human Genetics, Quantitative Biology Institute, and Institute for Computational Health Sciences, University of California, San Francisco, California, USA.; Chan-Zuckerberg Biohub, San Francisco, California, USA.

## Abstract

The potential impact of structural variants includes not only the duplication or deletion of coding sequences, but also the perturbation of non-coding DNA regulatory elements and structural chromatin features, including topological domains (TADs). Structural variants disrupting TAD boundaries have been implicated both in cancer and developmental disease; this likely occurs via ‘enhancer hijacking’, whereby removal of the TAD boundary exposes enhancers to new target transcription start sites (TSSs). With this functional role, we hypothesized that boundaries would display evidence for negative selection. Here we demonstrate that the chromatin landscape constrains structural variation both within healthy humans and across primate evolution. In contrast, in patients with developmental delay, variants occur remarkably uniformly across genomic features, suggesting a potentially broad role for enhancer hijacking in human disease.

---

The prevalence and potential impact of structural variants is increasingly appreciated (Cheng et al., 2005; Chiang et al., 2017; Zhang and Lupski, 2015). In addition to disrupting coding sequences through deletion, duplication, or inversion, structural variants can perturb the relative arrangement of non-coding DNA regulatory elements and structural features of the chromatin landscape, with consequences in development and disease (Krijger and de Laat, 2016; Spielmann and Mundlos, 2016). Chromatin boundaries at the borders of topologically associating domains (TADs (Dixon et al., 2012; Nora et al., 2012)), largely dependent on the binding of CTCF (Nora et al., 2017; Wutz et al., 2017), are structural features of the genome of much recent interest and are hypothesized to play an important role in gene regulation.

An emerging line of research implicates structural variants as functionally significant by altering TAD boundaries in cancer (Akdemir et al., 2017; Hnisz et al., 2016; Wala et al., 2017; Weischenfeldt et al., 2017). One likely mechanism is ‘enhancer hijacking’ (Beroukhim et al., 2016; Northcott et al., 2014), also previously termed ‘enhancer adoption’ (Lettice et al., 2011), whereby a structural variant removes or moves a TAD boundary to expose TSSs to regulatory enhancers from which they would normally be insulated. While there have been intriguing examples of TAD boundary disruptions in developmental diseases (Franke et al., 2016; Kraft et al., 2015; Lupiáñez et al., 2015; Symmons et al., 2016), the effect of structural variants on chromatin features like TAD boundaries has received relatively little systematic attention outside of cancer (Ibn-Salem et al., 2014), until the past year (Flöttmann et al., 2017; Huynh and Hormozdiari, 2018; Krefting et al., 2017; Lazar et al., 2017; Redin et al., 2017; Zepeda-Mendoza et al., 2017).

To systematically test if TAD boundary disruptions are under negative selection and compare their evolutionary constraint to that of other regulatory elements, we examined patterns of structural variation across evolutionary time scales from fixed differences between ape genomes to rare variants in human populations (**Fig. 1**). This strategy is motivated by the fact that the ability of negative selection to purge a given variant from the population will depend on how deleterious it is and how much time selection has had to act on it (Hartl and Clark, 1998). Hence, we can infer relative levels of functional constraint on TAD boundaries by comparing the frequency with which they are altered by structural variants across evolutionary time scales and comparing this frequency with that of other genomic elements and chromatin states, such as transcription start sites, enhancers, and heterochromatic regions. We find that deletions are strongly depleted at active chromatin states and TAD boundaries. This signature of negative selection is strikingly absent in patients with autism and developmental delay, where deletions occur remarkably uniformly across the genome. Together our analyses uncover a genome-wide pattern of negative selection against structural variants that would have the potential to alter chromatin structure and lead to enhancer hijacking.

**Figure 1:**
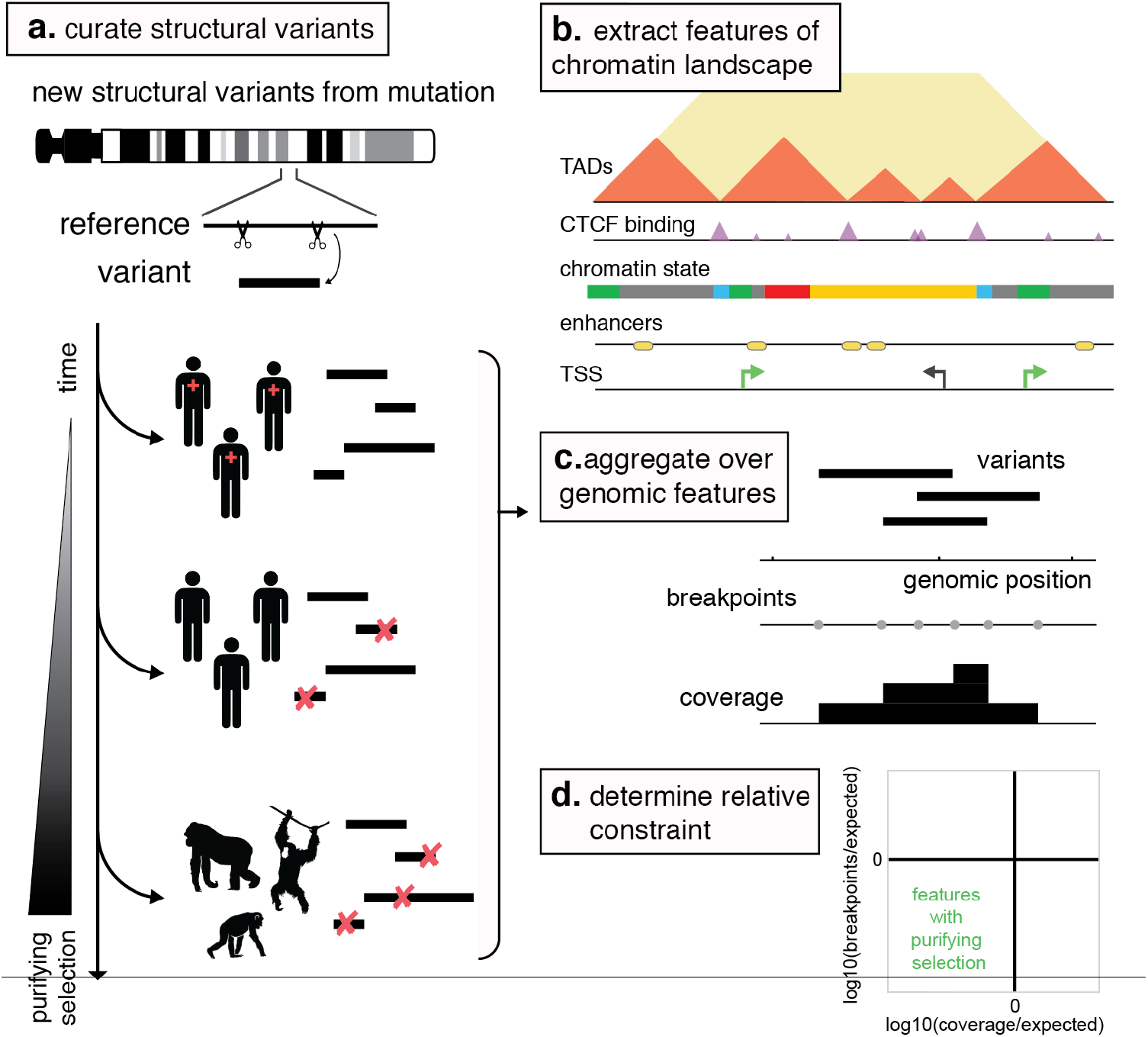
Approach to detect purifying selection against deleterious structural variants across the human genome over evolutionary timescales. **A.** To study sets of structural variants that have been subject to purifying selection for decreasing amounts of time, we obtained structural variants representing divergence with great apes (Sudmant et al., 2013), variation within the human population (Coe et al., 2014), and those detected in patients (shown with red crosses) with developmental delay and autism (Coe et al., 2014). **B**. To characterize the chromatin landscape, we curated the following genomic features: chromatin states from ENCODE Roadmap (Roadmap Epigenomics Consortium et al., 2015), cross-tissue gene expression from GTEx (GTEx Consortium, 2015), CTCF binding clusters from ENCODE (ENCODE Project Consortium, 2012) and TAD boundaries from high-resolution Hi-C data (Rao et al., 2014). **C**. To assess the distribution of structural variants across the human genome, we summarize each set as: 1) the frequency of breakpoints, and 2) their coverage across the genome. **D**. We then determine whether genomic features are enriched or depleted for variant breakpoints and coverage. As structural variants subject to purifying selection are gradually removed from the population and hence become less common over increasing evolutionary timescales, we expect features under purifying selection to be depleted for breakpoint frequency and coverage.

## Results

### Data and Methods

To study structural variants that have been subject to selection for different periods of time, we obtained sets representing divergence with great apes (Sudmant et al., 2013), variation within the human population (Coe et al., 2014), and those detected in patients with developmental delay and autism (Coe et al., 2014). To assess their genome-wide impact, we summarize each set of structural variants as: 1) the frequency of breakpoints, and 2) their coverage across the genome (**Fig. 1**). Breakpoints capture how likely a variant is to start or stop at a particular genomic position, whereas coverage represents the number of variants that have altered a particular genomic position. While related, these could in principle capture different factors; for example, a key genomic feature could be adjacent to a region prone to frequent breaks, yet be locally depleted for deletions that remove it. To enable analyses relating to the frequency of a variant in the population, we used unique combinations of start and end points to determine shared variants. We focus on deletions, as duplications can either be in tandem, adjacent to the original copy, or elsewhere in the genome, adding additional complexity to interpreting their effects (Ibn-Salem et al., 2014).

To characterize the chromatin landscape, we curated the following genomic features: chromHMM chromatin states from Roadmap (Roadmap Epigenomics Consortium et al., 2015), cross-tissue gene expression for TSSs from GTEx (GTEx Consortium, 2015), TAD boundaries from high-resolution Hi-C data, called using an arrowhead score (Rao et al., 2014), and CTCF binding clusters from ENCODE (ENCODE Project Consortium, 2012). CTCF frequently demarcates TAD boundaries and CTCF ChIP-seq data is currently available for a broader set of cell-types than is high-resolution Hi-C data. We quantified the strength of a TSS in GTEx as the sum of its expression across human tissues, because genetic variants have potential to alter function in any tissue. Similarly, we quantified the strength of a CTCF cluster as its aggregate binding across cell lines. TSSs and the midpoints of CTCF clusters were extended +/-5kb to enable consistent comparisons with TAD boundaries.

To quantitatively evaluate relative levels of purifying selection on the genomic features defined above, it is critical to appropriately normalize deletion rates by their expected levels. We quantified this expectation as a uniform distribution across the genome, given the proportion of the genome covered by that genomic feature (**Methods**). By similarly computing observed versus expected rates for other classes of genomic elements, the relative levels of purifying selection at TAD boundaries can be quantitatively evaluated. We refer to a genomic feature with fewer variants than expected as “depleted”. Our organizing principle is that the relative depletion of variants for a given genomic feature represents the average functional importance of that feature.

### Ape deletions are strongly depleted at active chromatin states

We first investigated the relationship between great ape deletions and human chromatin states. We considered deletions relative to the human genome that were fixed in at least one ape species (Bornean and Sumatran orangutans, any of four chimpanzee subspecies, bonobos, and Eastern and Western gorillas (Sudmant et al., 2013)), additionally requiring that these deletions were parsimonious (i.e. not better explained by duplication in the human lineage). We found that both deletion breakpoints and coverage were depleted in active chromatin states (**Fig. 2A**), consistent with purifying selection acting to purge deletions affecting transcriptionally active portions of the genome. Indeed, only quiescent chromatin and heterochromatin were not consistently depleted for either coverage or deletion breakpoints across cell-types. TAD boundaries were also avoided by deletions, and avoided slightly more on average than TSSs. Confirming these observations, we found similar patterns for a more recently characterized set of gorilla deletions (Gordon et al., 2016)(**Supplemental Fig. 1**).

**Figure 2:**
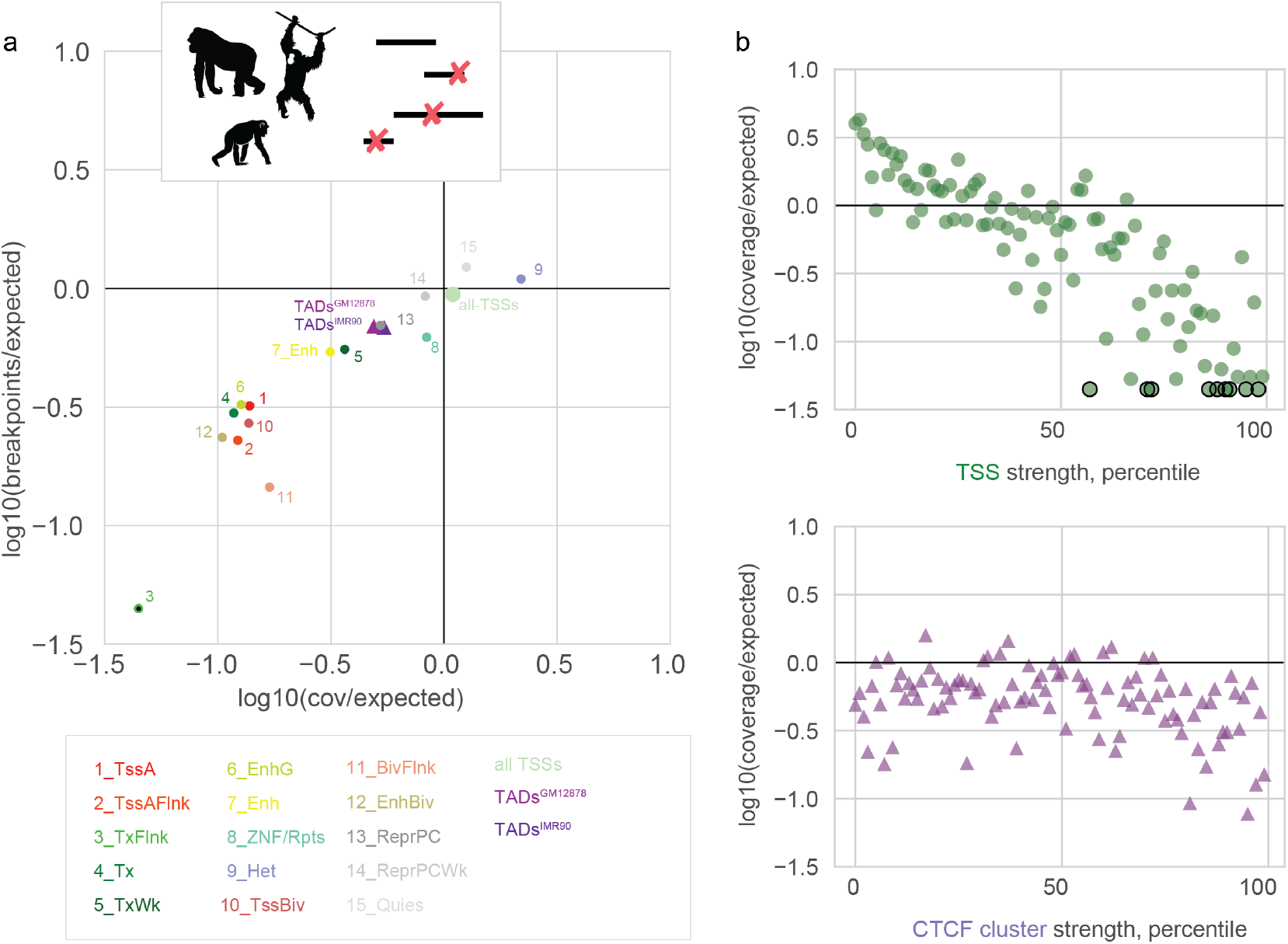
Ape deletions show patterns of purifying selection at active chromatin states, CTCF clusters, and TAD boundaries. **A.** Deletions observed in apes have both lower coverage and breakpoint frequency than expected in active genomic features and at TAD boundaries. Crosses show the 25^th^ and 75^th^ percentiles across Encode Roadmap cell types. A black endpoint indicates that no variants were observed for that chromatin class, and the corresponding bar was truncated for display. **B.** Ape deletion coverage at TSSs (green circles, top) and CTCF clusters (purple triangles, bottom) scales with the strength of these genomic features. Each point represents the average over one of 100 quantiles; black edges indicate quantiles with no observed deletion coverage shown at the minimal plotted y-value, for display.

We next examined if the strength of negative selection at TSSs and CTCF binding clusters relates to the strength of these features. Indeed, coverage was more depleted for TSSs that were more highly expressed (**Fig. 2B**), consistent with stronger purifying selection acting on deletions at more broadly important genes. Similarly, we found that both breakpoints and coverage were more depleted for CTCF clusters that were more strongly bound in aggregate (**Fig. 2B**). Interestingly, CTCF clusters were more avoided than TSSs up to ~50^th^ percentile of aggregate GTEx expression. Collectively these findings are indicative of purifying selection acting to remove deleterious variants that would perturb functionally important chromatin features, including TAD boundaries, at the timescale of great ape evolution.

### Human deletions reveal the spectrum of selective constraint at chromatin features

We next investigated the connection between deletions found in healthy humans (Coe et al., 2014) and chromatin features (**Fig. 3**). These include structural variants that are segregating in the human population and hence have not been under selection for as long as deletions that are fixed differences between apes. Nonetheless, we expect deleterious structural variants should be depleted in healthy adults. As observed for apes, human deletions were depleted in active chromatin states and at TAD boundaries (**Fig. 3A**), again arguing for purifying selection acting to purge deletions that impact these chromatin features. We found similar, though less pronounced, patterns (**Supplemental Fig. 1**) across chromatin states in an independent set of human deletions from a smaller set of individuals (Sudmant et al., 2015), and note that a similar avoidance of TAD boundaries was reported for International Cancer Genome Consortium (ICGC) germline deletions (Akdemir et al., 2017). As observed for ape deletions, TSSs were more strongly depleted if more highly expressed and CTCF clusters were more strongly depleted if more strongly bound (**Fig. 3B**); these consistent relationships for ape variants and healthy human variants argue that the strength of purifying selection is directly related to the importance of a chromatin feature. Additionally, CTCF clusters were more avoided than TSS up to the ~60^th^ percentile of aggregate GTEx expression, suggesting that these non-coding features could be as important as many coding features. Interestingly, CTCF motifs alone are not particularly depleted (**Supplemental Fig. 2A**), even after stratifying by the quality of the motif (Grant et al., 2011), consistent with only a fraction being sufficiently occupied to enact their functional roles (Nora et al., 2017). Together, these findings argue that purifying selection acts to remove deleterious variants that would perturb functionally important chromatin features within the human population.

**Figure 3:**
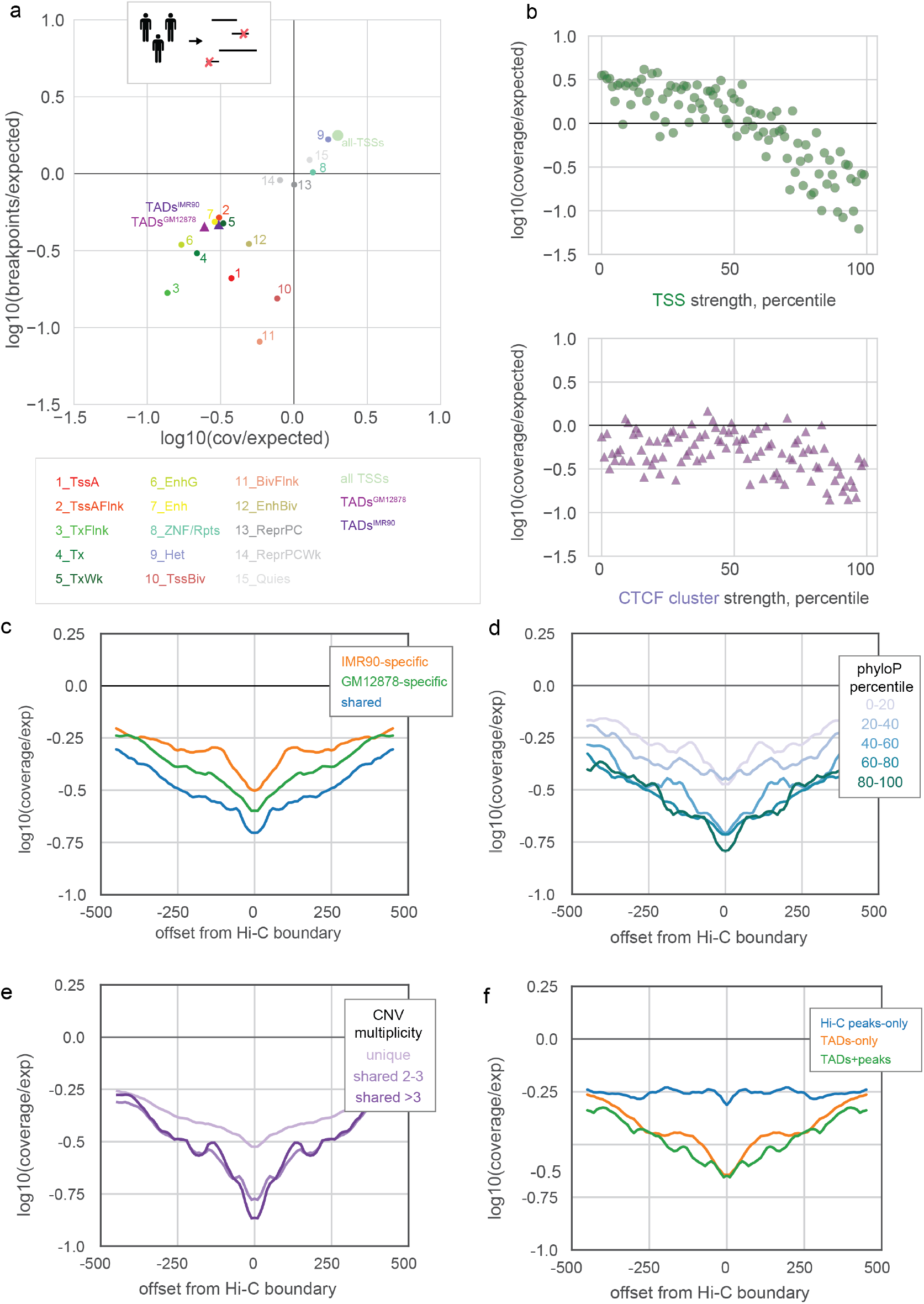
Human deletions reveal the spectrum of purifying selection across genomic features. **A**. Deletions observed in healthy humans have lower coverage and breakpoint frequency in active states and at TAD boundaries. Crosses show the 25^th^ and 75^th^ percentiles across Encode Roadmap cell types. A black endpoint indicates that no variants were observed for that chromatin class, and the corresponding bar was truncated for display (as in **Fig. 2A**). **B**. Healthy human deletion coverage at TSSs (green circles, top) and CTCF clusters (purple triangles, bottom) scales with the strength of these genomic features. Each point represents the average over one of 100 quantiles. **C**. TAD boundaries shared across cell types are more depleted for human deletions than those found in only one cell-type, shown by coverage in the +/-500kb genomic region at 10kb binned resolution. **D**. TAD boundaries with more evolutionary conservation at the base-wise level are also more depleted for human deletions, as shown by coverage in the +/-500kb genomic region around TAD domain boundaries, stratified by their average phyloP. **E**. Human deletions that are shared across individuals are more depleted at TAD boundaries, as shown by coverage in the +/-500kb genomic region around all GM12878 TAD boundaries. **F**. TAD boundaries are more depleted for human deletions than Hi-C peaks, as shown by coverage in the +/-500kb genomic region around these features.

Leveraging the larger number of deletions in this dataset, we next investigated the coverage of deletions not only at TAD boundaries, but in the surrounding region as well (**Fig. 3C-F**). This analysis reveals that deletions are broadly depleted around TAD boundaries, and most depleted right at boundary sites. Using this approach, we additionally find that (i) boundaries called in multiple cell types are more depleted (~1.4- fold for two versus only one cell-type, **Fig. 3C**); (ii) boundaries with higher average base-wise conservation (phyloP score, (Pollard et al., 2010)) are more depleted (~2.2-fold more for the top versus bottom quintile, **Fig. 3D**); and (iii) that deletions present in multiple people are more depleted at boundaries (~1.7-fold, **Fig. 3E**), consistent with shared variants having spent more time under purifying selection.

Surprisingly, we found that avoidance of deletions at boundary elements shows little dependence on boundary strength measured by within cell-type insulation (**Supplemental Fig. 2B, and Methods**), suggesting that the called set of boundaries all provide sufficient insulation to regulate genes in their neighborhoods. As boundaries are thought to play roles in transcriptional insulation and might present the greatest risk for enhancer hijacking when insulating genes with very different expression, we tested if boundaries over which GTEx expression is discordant show stronger signatures of deletion avoidance. Aggregating gene expression in a window on each side of a boundary within each tissue-type and taking the maximal difference between the two sides across tissue-types, we only found limited evidence in support of this hypothesis (**Supplemental Fig. 2C**).

Since TSSs of active genes are avoided by deletions, we next sought to test if the observed depletion at TAD boundaries could be a consequence of their genomic proximity to actively expressed genes. When we stratified boundaries by their distance to the nearest highly-expressed TSS, however, we saw that the depletion leveled out to average levels after ~100kb (**Supplemental Fig. 2D**). This result argues that purifying selection can act directly to remove deleterious variants that would perturb TAD boundaries.

Another commonly discussed feature of chromosome folding is the focal point-to-point peaks of contact frequency observed in Hi-C maps, associated with strong CTCF binding sites overlying oriented motifs (often termed loops, (Rao et al., 2014)). When we subjected this set of regions to the same analysis, we found that TAD boundaries are more depleted than are the bases of Hi-C peaks (~2.2-fold, **Fig. 3F**). Consistently, we found boundaries are also more conserved at the single-nucleotide level than are peak bases, as measured by either their maximum or average phyloP score (**Supplemental Fig. 2E**). Together this argues for TAD boundaries generally having either broader, or more important, functional roles than Hi-C peaks.

### Active chromatin states and chromatin boundaries are disrupted in patients with developmental delay or autism

To investigate the situation where purifying selection has had the least time to act, we considered the pattern of deletions in patients with developmental delay or autism. In contrast with deletions from apes and healthy humans, deletions in affected individuals displayed no avoidance of TSSs or CTCF clusters, regardless of the strength of these genomic features (**Fig. 4A-D**). Consistently, active chromatin states and TAD boundaries were also not avoided by this set of deletions (**Supplemental Fig. 1, Supplemental Fig. 2F**), whereas they were in controls from the same study.

**Figure 4:**
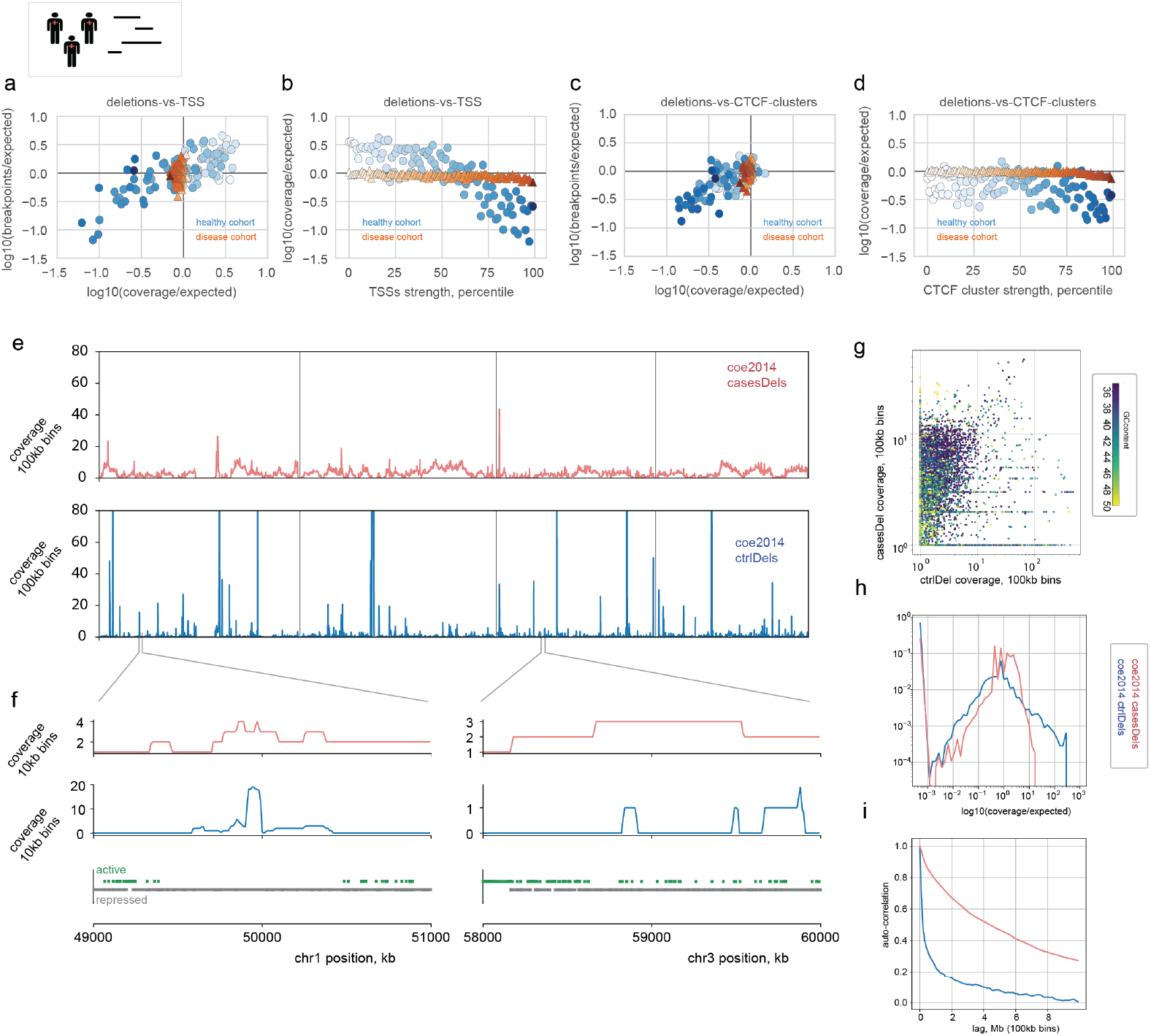
Deletions in human disease show no avoidance of key genomic features. **A**. Unlike for healthy humans, deletions in patients with developmental delay and autism show no avoidance of strong TSSs, either in terms of their coverage, or their breakpoint frequency. Each point represents average deletion coverage and breakpoint frequency for TSSs stratified by and shaded by strength, for variants found in healthy humans (blue) and variants found in patients (orange). **B**. Deletion coverage in patients shows little relationship with TSS strength. **C-D.** As for A-B, but for CTCF clusters, stratified by strength. **E**. 100kb binned coverage profiles of deletions from patients (cases, red) and healthy controls (blue) across the first four chromosomes illustrate differences in their large-scale distribution across the genome. **F**. zoom into 10kb binned coverage profiles of deletions for the indicated genomic regions, above tracks showing inactive (grey, 8_ZNF/Rpts, 9_Het, 13_ReprPC, 14_PeprPCWk, 15_Quies) versus active (green, other states) Roadmap states in these regions. The region on chr1 *(left)* shows an island of high coverage in controls over a broad heterochromatic state; the region on chr3 *(right)* shows broadly elevated coverage in cases, as compared with the more punctuated coverage in controls. **G**. scatter plot of deletion coverage, colored by GC content, showing a low correlation of coverage profiles at the 100kb level between cases and controls. **H**. distribution of coverage per 100kb bin showing a rapid decay for controls and a more gradual decay in cases. **I**. autocorrelation of 100kb binned profiles of deletion coverage, showing longer autocorrelation length and more slowly varying coverage profiles in patients.

In fact, deletions in patients display a remarkably uniform distribution across the genome (**Fig. 4E-I**), in addition to being longer (Coe et al., 2014), as compared with deletions in healthy individuals. This is observed for deletions in patients both in the more slowly decreasing autocorrelation (**Fig. 4I**) and the less skewed distribution (**Fig. 4H**) of the coverage profiles. We also note that the coverage profile of deletions in patients is not particularly correlated with that in controls (**Fig. 4G**).

Given that deletions are not depleted at boundaries in patients with developmental disease or autism, an interesting possibility is that deletions of specific TAD boundaries could broadly contribute to disease etiologies beyond cancer. To address this, we permuted positions of deletions, calculated coverage profiles of these permuted events, and used these profiles to determine a threshold for significantly deleted 10kb regions separately for cases and controls, using the 99.9^th^ percentile of the respective permuted coverage profile. We found that significantly deleted 10kb regions were enriched at boundaries in cases, relative to controls (Fisher’s exact test, OR 1.47, p-value <1e-4, **Table 1A**).

**Table 1.**
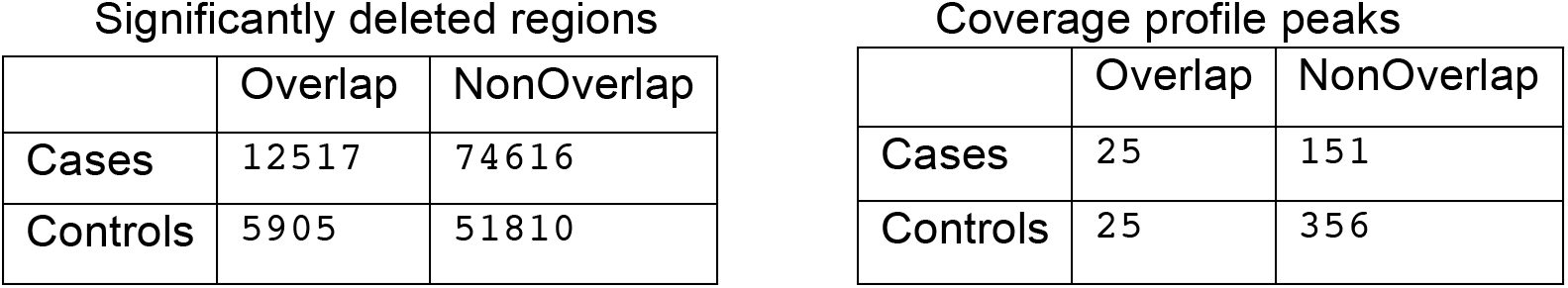
**A**. Table for significantly deleted regions in cases and controls, and whether they overlap TAD boundaries. **B**. Table for peaks in 10kb binned coverage profiles in cases and controls, and whether they overlap TAD boundaries.

|  | Overlap | NonOverlap |
| --- | --- | --- |
| Cases | 12517 | 74616 |
| Controls | 5905 | 51810 |

|  | Overlap | NonOverlap |
| --- | --- | --- |
| Cases | 25 | 151 |
| Controls | 25 | 356 |

To determine possible functional roles of deleted boundaries, we considered the enrichment of gene ontology categories for genes around TAD boundaries that were significantly deleted for the ape, healthy human, and developmental disease deletions using GO-rilla (Eden et al., 2009). Interestingly, these three gene sets displayed different GO term enrichments: ape deletions had terms related to sensory perception; healthy humans had immune-related terms; and developmental disease deletions had chromatin-related terms (**Supplemental Table 1**). We note that these results for genes near significantly deleted TAD boundaries in apes are in agreement with gene-based approaches that report recurrent deletions in olfactory perception loci across apes (Dong et al., 2009).

We then reasoned that local maxima, or peaks, in the genome-wide deletion coverage profile that overlap particular TAD boundaries could strengthen the case for a given boundary’s putatively causal role in disease. A similar approach has been used for implicating particular genes from somatic copy alterations in cancer (Beroukhim et al., 2010). We found that peaks in the coverage profiles were moderately enriched (OR 2.36, p-value .00610, **Table 1B**). Since this genome-wide enrichment was relatively mild, we refrained from determining the significance of individual boundary elements in this patient cohort. Indeed, a challenge of using patient deletions to determine the role of individual TAD boundaries is that deletions in the disease cohort are particularly large (Coe et al., 2014), making it difficult to ascribe a role that primarily relates to disrupting the integrity of 3D genomic folding.

Nevertheless, by visual inspection we note there are intriguing candidates for future analyses, including a highly focally deleted boundary on chromosome 18 that appears to insulate a TAD containing the RNA-binding protein MEX3C from an adjacent TAD containing the gene DCC, involved in neurogenesis (**Fig. 5**). Combined with our observations that disruptions to TAD boundaries are generally avoided in healthy human cohorts, these results indicate that disruption of TAD boundaries could play important roles in diseases outside of those established in cancer.

**Figure 5.**
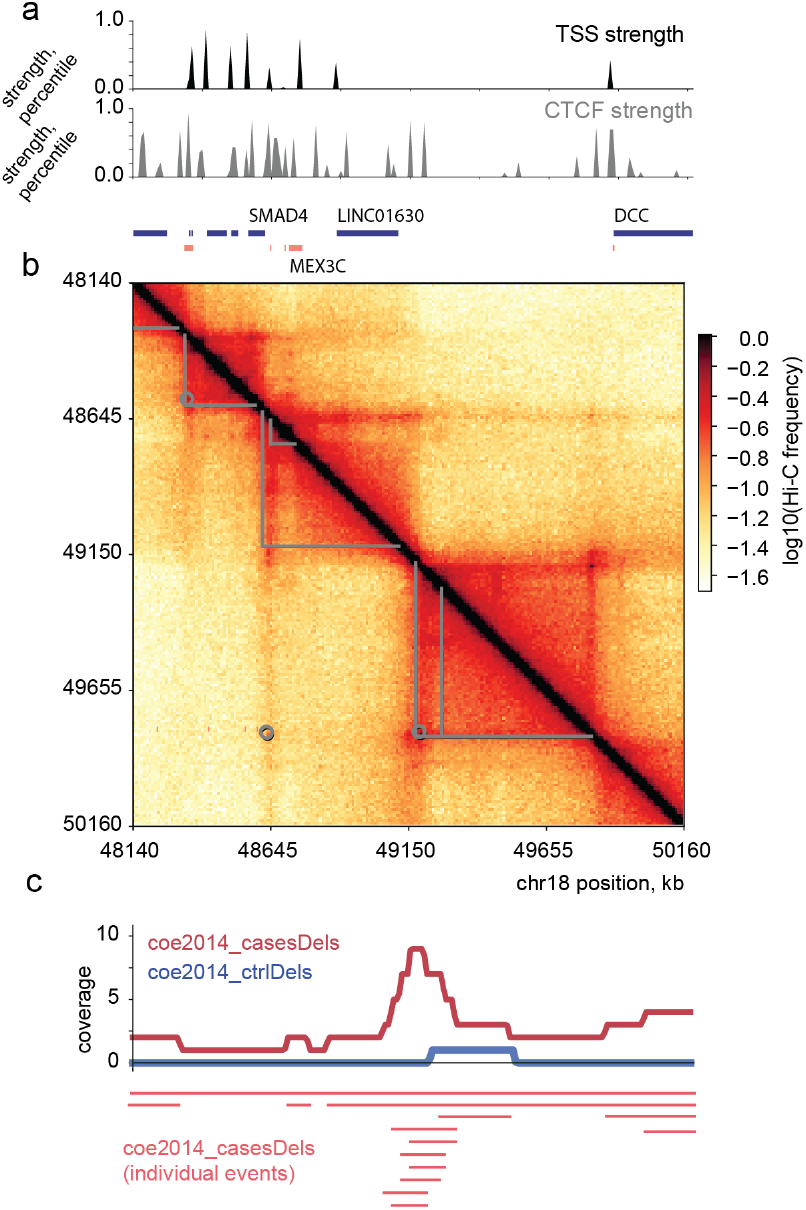
Focal enrichment of deletions in cases at a TAD boundary on chr18. **J**: *(top)* 10kb binned profiles of TSS and CTCF cluster strength in this region, *(bottom)* positions of genes colored by orientation (blue, forward; red, reverse). **K**: Hi-C map for this region from GM12878 cells at 10kb resolution (Rao et al., 2014), with associated TAD and Hi-C peak calls overlaid as grey lines and circles. **L**: Coverage of deletions in patients (cases, red) and controls (blue) over this region; red bars below show individual events in patients that build up this coverage profile.

### Duplications display a more complex relationship with chromatin features than deletions

We next considered how functional constraint influences the patterns of both duplications and deletions across evolutionary time scales. For a given level of average constraint on a class of genomic features, we expect structural variants to be most avoided for apes, then healthy humans, followed by humans with diseases, reflecting decreasing time for selection to have operated. This is indeed what we observe for deletions of TSSs, as would be expected if they were generally deleterious and under purifying selection (**Supplemental Fig. 3A**). Unexpectedly, CTCF clusters seem to be similarly, or even slightly less, avoided for deletions in healthy humans and apes (**Supplemental Fig. 3B**). For healthy humans, we observed similar, yet less pronounced, patterns for duplications than for deletions. Interestingly, longer duplications were the main contributor to the avoidance of more active TSSs and strongly bound CTCF sites observed for healthy human variants (**Supplemental Fig. 4**). This may indicate a greater importance of genomic context for duplications, which may be one important factor for understanding the observed differences between these classes of structural variants. Surprisingly, ape duplications showed no clear trend for TSSs or CTCF clusters, which held after stratifying by length (**Supplemental Fig. 4**), in contrast to duplications in healthy humans. However, we note that ape duplications are on average much shorter than those in healthy humans, and the shortest human duplications also show little avoidance of TSSs or CTCF clusters (**Supplemental Fig. 4**). As synteny breakpoints are avoided within TADs (Krefting et al., 2017; Lazar et al., 2017), our observations support the concept that the details of how a given structural variant impacts genomic organization determines its effect on fitness.

## Discussion

In summary, we find evidence for purifying selection acting on structural variants, depending on their local chromatin context. Not only are deletions depleted in active chromatin states both in apes and the human population, but also at CTCF sites and TAD boundaries. Indeed, boundaries are avoided as strongly as intermediately expressed TSSs, suggesting parts of the coding and non-coding genome could be equally important from the point of view of deletions. In contrast with these sets of variants that had time to experience purifying selection, we found that variants present in patients with autism and developmental delay were surprisingly uniform across chromatin states, and displayed no preferential avoidance of strongly expressed TSSs or strongly bound CTCF sites.

Interestingly, the uniformity across features we observed here for autism and developmental delay deletions contrasts with recent observations of cancer deletions, where deletion frequency has been reported as closely related to chromatin state (Wala et al., 2017) for the Pan-Cancer Analysis of Whole Genomes (PCAWG) dataset. Indeed, when we analyzed cancer deletions from COSMIC with the same methods used above, we found a very different pattern from that in developmental delay patients, and actually opposite that found in healthy individuals (**Supplemental Fig. 5**). We speculate this may either stem from different mutational mechanisms for somatic alterations in cancers as compared with deletions in autism and developmental delay patients, including transcription-related mutagenesis for deletions in cancer, or widespread positive selection for deletions in cancer genomes.

Our findings further argue that structural variants with the potential to lead to enhancer hijacking are under purifying selection. Interestingly, the overall distribution of both deletions and duplications in healthy humans rapidly plummets after ~2Mb (Coe et al., 2014), which is also roughly the furthest distance over which enhancers are known to act (Krijger and de Laat, 2016), the size of the largest TADs (Bonev and Cavalli, 2016), and the distance over which cohesin enriches contact frequency (Fudenberg et al., 2018).

Put another way, it appears that deletions or duplications bringing genomic elements together that would otherwise never communicate are particularly avoided. Further supporting the idea that enhancer hijacking is imperative to avoid, we note that very broadly expressed genes tend to be closer to very broadly bound CTCF sites (**Supplemental Fig. 2G**), consistent with a fundamental role of CTCF in constraining ectopic expression (Ing-Simmons et al., 2015; Willi et al., 2017).

Our results are also consistent with emerging mechanistic insights into enhancer-promoter communication (Dekker and Mirny, 2016). Our finding that Hi-C peaks are less avoided by deletions than TAD boundaries raises the possibility that TAD boundaries may generally have either broader, or more important, functional roles than Hi-C peaks.

If enhancer-promoter contacts are very dynamic (Fukaya et al., 2016; Gu et al., 2018) and enhancers are promiscuous (de Laat and Duboule, 2013) it may be relatively more important to keep enhancers from ectopically activating genes rather than specifying very specific enhancer-promoter pairings. Alternately, boundaries may be more important if they are more stable across cell-types, and orchestrate different sets of peaks in different cell types.

An important caveat for using structural variants to assay functional importance of different genomic regions is the non-uniformity of genome. Indeed, active regulatory elements are clustered along the genome (ENCODE Project Consortium, 2012; Filion et al., 2010), making it difficult to discern their independent casual roles from a set of structural variants that can span multiple genomic features. Nevertheless, this property of structural variants can be beneficial for characterizing the chromatin landscape if disruptions of multiple elements, e.g. bound CTCF sites, are required to alter the boundary activity of TADs, as appears to be the case at the *HoxD* locus (Rodríguez-Carballo et al., 2017).

Collectively, our findings that TAD boundaries and strong CTCF sites are more important than many low-expressed coding sequences argue for re-thinking the gene-centric paradigm for interpreting structural variants. Our results also support efforts to broadly characterize the epigenome, beyond assaying transcription, as many non-coding regions are more avoided by structural variants than annotated TSSs with low expression levels.

## Online Methods

### Structural variant datasets

We focused our analysis on deletions and duplications from (Sudmant et al., 2013) and (Coe et al., 2014). (Coe et al., 2014) variant calls were obtained from https://www.ncbi.nlm.nih.gov/dbvar/studies/nstd100/, supporting_variants_for_nstd100_coe2014.csv).

We used liftOver to convert variants from (Sudmant et al., 2013) from hg18 to hg19 coordinates and COSMIC variants (http://cancer.sanger.ac.uk/cosmic/download, CosmicCompleteCNA.tsv.gz, release v84) from hg38 to hg19; all other datasets had hg19 coordinates available. We also analyzed variants from (Gordon et al., 2016) and (Sudmant et al., 2015). To enable analyses relating to the frequency of a variant in the population, we used unique combinations of start and end points to determine shared variants. We limited all analyses to autosomes.

### Chromatin and expression datasets

Chromatin state analyses were performed using the core 15-state model across 127 cell types from Roadmap ((Roadmap Epigenomics Consortium et al., 2015), http://egg2.wustl.edu/roadmap/web_portal/chr_state_learning.html).TSS analyses were performed using GTEx v6 release ((GTEx Consortium, 2015), https://www.gtexportal.org/home/), using GTEx_Analysis_v6p_RNA-seq_RNA-SeQCv1.1.8_gene_median_rpkm.gct.gz for expression values and encode.v19.genes.v6p_model.patched_contigs.gtf.gz for TSS positions. From the above files, we quantified the strength of a TSS in GTEx as the sum of its expression across tissues. TAD boundary and Hi-C peak analyses were performed using arrowhead domains and hiccups loop lists from (Rao et al., 2014) downloaded from (https://www.ncbi.nlm.nih.gov/geo/query/acc.cgi?acc=GSE63525). CTCF binding clusters were obtained by downloading narrowPeak files from ENCODE (ENCODE Project Consortium, 2012) http://hgdownload.soe.ucsc.edu/goldenPath/hg19/encodeDCC/wgEncodeAwgTfbsUniform/ for the Broad center, and then using bedtools cluster on the aggregated set with a merge distance of 5kb. We quantified the strength of a CTCF cluster as its aggregate binding across samples. TSSs and the midpoints of CTCF clusters were extended +/-5kb to enable consistent comparisons with TAD boundaries.

### Relative abundance of structural variants

We quantified the relative abundance of variants for a genomic feature, displayed as the log10(observed/expected). For breakpoint frequency, the observed/expected was calculated as: 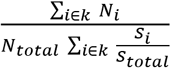, where *i* indexes genomic regions, S is the size of and N is the number of variant breakpoints in a given region *i*, summed over regions in the class *k* (e.g. chromatin state, or quantile of CTCF binding strength). Intersection of variant positions and genomic features were performed using bedtools (Quinlan and Hall, 2010). The observed/expected for coverage was calculated similarly, except with N as the total number of basepairs covered by variants in a region *i* of class *k*. Variants starting in the first or last 2Mb of a chromosome, or within 2Mb of centromeric regions (defined by UCSC hg19 gap file) were excluded from analysis, as these may be more prone to artifacts (Coe et al., 2014). These regions were similarly excluded from the calculation of the total genome size, *S_total_*.

### Permutation analysis for boundary deletions

To generate coverage profiles for permuted deletions, we used bedtools shuffle using the hg19 autosomes as the genome, and the same excluded regions for the enrichment analyses above. For each set of variants we then took the 99.9^th^ percentile of the coverage profile as a threshold to identify significantly deleted regions. We then tabulated the number of significantly deleted regions that overlapped TAD boundaries from GM12878 data versus those that were in other places in the genome and performed a Fisher’s exact test on the resulting 2×2 table. To identify peaks in the coverage profiles we used the peakdet algorithm (Billauer E (2012). peakdet: Peak detection using MATLAB, http://billauer.co.il/peakdet.html), where the minimum required prominence is the same variant set specific threshold calculated above. We considered a coverage profile peaks as intersecting a TAD boundary if it was within +/- 10kb (i.e. one bin) from genomic location of the TAD boundary.

### Enrichment of GO terms for genes around TAD boundaries

To quantify the enrichment of GO terms around significantly deleted TAD boundaries, we used GO-rilla (Eden et al., 2009) a web-based application (http://cbl-gorilla.cs.technion.ac.il/) that can calculates enrichment both for ranked lists (Eden et al., 2007) and for a set of target genes versus a background set. We determined significantly deleted boundaries as TAD boundaries from GM12878 (Rao et al., 2014) with an observed coverage by deletions that exceeded the 99.9^th^ percentile a permuted coverage profile, calculated separately using 1000 permutations for each set of deletions. We then took all TSSs with non-zero GTEx expression +/- 500kb around each significantly deleted boundary as the three different target sets, and the background set as all TSSs +/- 500kb from any boundary (for lists of GO-rilla inputs **Supplemental Table 2**). We note that GO-rilla has annotations for only 45% of the TSSs on these input lists, as many gencode-V6 TSSs are un-annotated or non-protein-coding transcripts.

## Acknowledgements

The authors thank Sean Whalen, Chris McFarland, and Kadir Akdemir for feedback, and Nezar Abdennur and Anton Goloborodko for suggesting computational methodology. The authors also thank their colleagues who provided feedback, both digitally on the previous version posted to bioRxiv and in person at the CSHL Systems Biology and Keystone Chromatin meetings. This work was supported by NIMH grant #MH109907, NHLBI grant #HL098179, and the San Simeon Fund.

## Author Contributions

GF and KP conceived of the study, GF analyzed the data, GF and KP wrote the paper.

## Competing Interests Statement

The authors declare no competing interests.

