## Supplementary Materials for "Chromatin features constrain structural variation across evolutionary timescales" (from another version of this preprint: version 1, 2018-03-19)

**Contents:**

**Supplemental Fig. 1.**
**Supplemental Fig. 2.**
**Supplemental Fig. 3.**
**Supplemental Fig. 4.**
**Supplemental Fig. 5.**
**Supplemental Table 1 caption.**
**Supplemental Table 2 caption.**

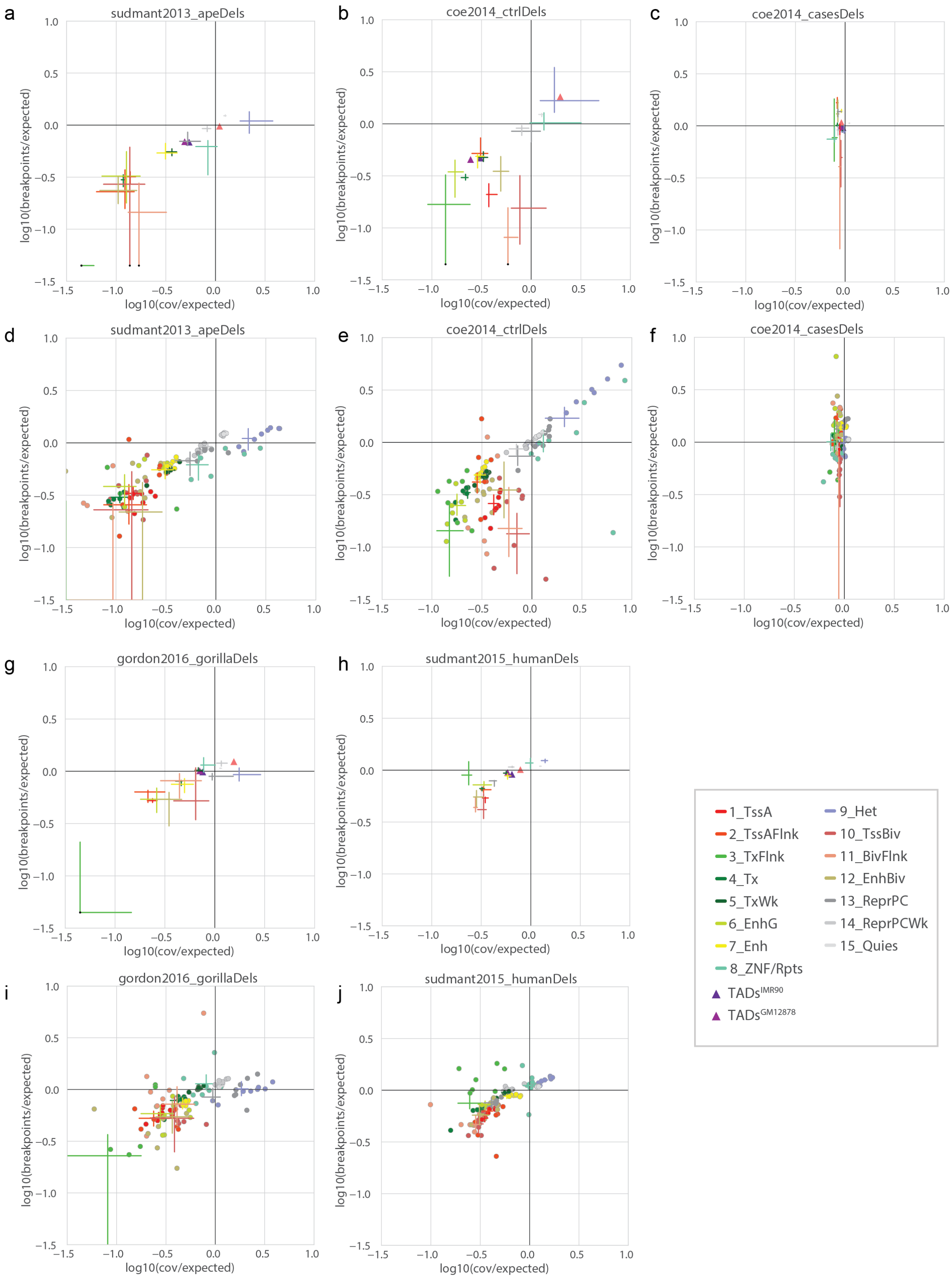


**Supplemental Fig. 1. (caption on next page)**

**Supplemental Fig. 1.**
**a-c, g-h.** Deletions observed in apes and healthy human have both lower coverage and breakpoint frequency than expected in active genomic features and at TAD boundaries. Crosses show the $25^{th}$ and $75^{th}$ percentiles across Encode Roadmap cell types. A black endpoint indicates that no variants were observed for that chromatin class in the $25^{th}$ percentile cell type, and the corresponding bar was truncated for display. Dataset for deletions is indicated above the associated plot. Note gorilla deletions (**g, i**) from (Gordon et al., 2016) show similar patterns to ape deletions (**a,d**). Also note human deletions from (Sudmant et al., 2015) (**h,j**) show similar, though less pronounced, patterns as compared with healthy humans from (Coe et al., 2014) (**b,e**).
**d-f, i-j.** Crosses as in other panels, points represent individual Roadmap ESC cell types.

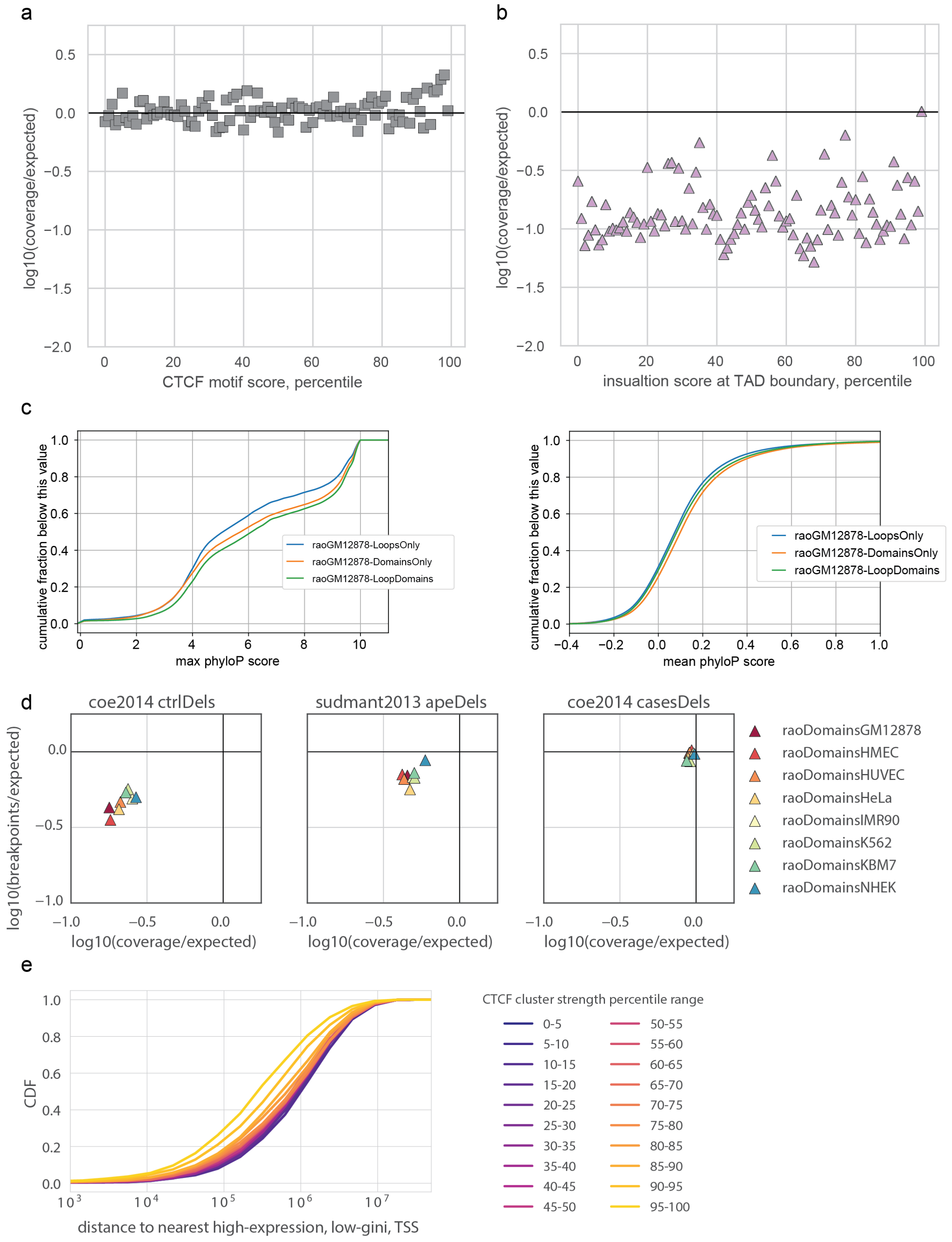


**Supplemental Fig. 2. (caption on next page)**

**Supplemental Fig. 2.**

**a.** Deletion coverage versus CTCF motif quality (Grant et al., 2011) for 100 quantiles.

**b.** Deletion coverage versus insulation score for TAD boundaries from GM12878 Hi-C, for 100 quantiles of insulation score. Insulation score is quantified by the contact frequency in a 250kb sliding window along the genome, as previously (Nora et al., 2017); note this within-cell type score differs greatly from the cross-cell-type measures of CTCF cluster strength and TSS strength used elsewhere.

**c.** ***Left:*** Cumulative distribution of maximum phyloP score in the 10kb window defined by the indicated feature; these three categories were obtained by intersecting the set of TAD domain boundaries and Hi-C peak bases from GM12878, and classifying a region as TAD-only (DomainsOnly), peak-only (LoopsOnly), or shared (LoopDomains). ***Right:*** Cumulative distributions of mean phyloP score in the same regions as above.

**d.** Deletion coverage versus breakpoint frequency for TAD boundaries across cell types for the indicated datasets. Healthy human and ape deletions are consistently depleted across cell types at TAD boundaries, patients with developmental delay and autism consistently show no depletion for either breakpoint frequency or coverage.

**e.** Cumulative distributions of the distance between a CTCF cluster and the nearest broadly and highly-expressed TSSs (below $10^{th}$ percentile Gini index, and above $90^{th}$ percentile aggregate expression), stratified by CTCF cluster strength (20 quantiles). Note that strongly bound CTCF sites across ENCODE cell types are closer to broadly- and highly-expressed TSSs characterized by GTEx, as indicated by the leftward shift of the CDF.

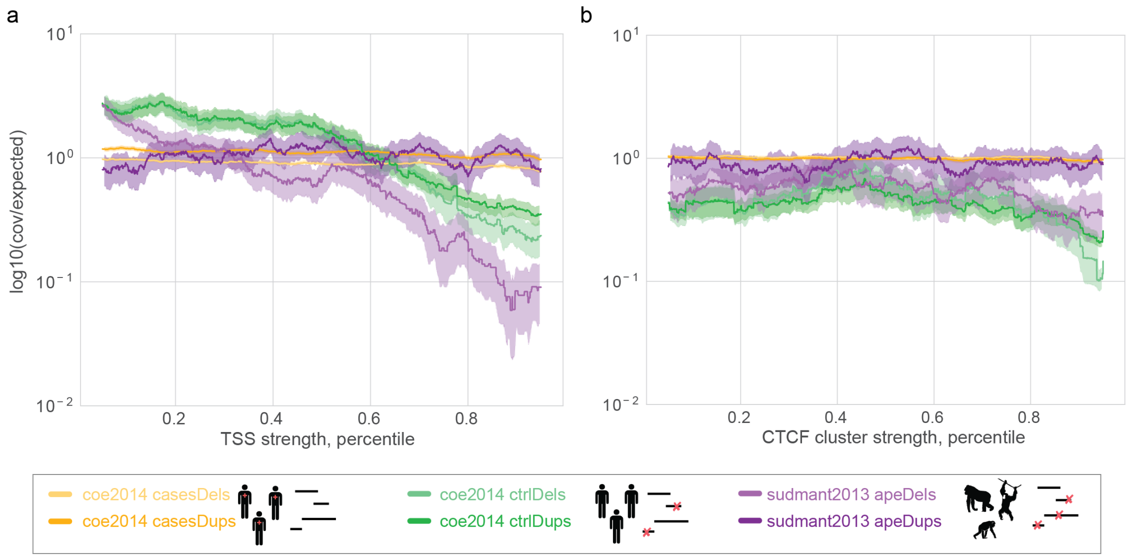


**Supplemental Fig. 3.**

**a.** For deletions, strong TSSs are most avoided for apes, then for healthy humans, and are not avoided in developmental delay patients, suggesting that these disease-associated deletions may be deleterious. Duplications show less avoidance of TSSs than deletions in healthy humans, and no avoidance for ape duplications. Curves show average expected coverage as a function of TSS strength in a sliding window; areas represent 5th & 95th percentiles of sliding mean calculated over 1000 bootstrap samples.

**b.** CTCF clusters are depleted for deletions across CTCF cluster strengths in both apes and healthy humans, consistent with their disruption being generally deleterious and under purifying selection. Similarly to TSSs, ape duplications display no preferential avoidance of CTCF clusters.

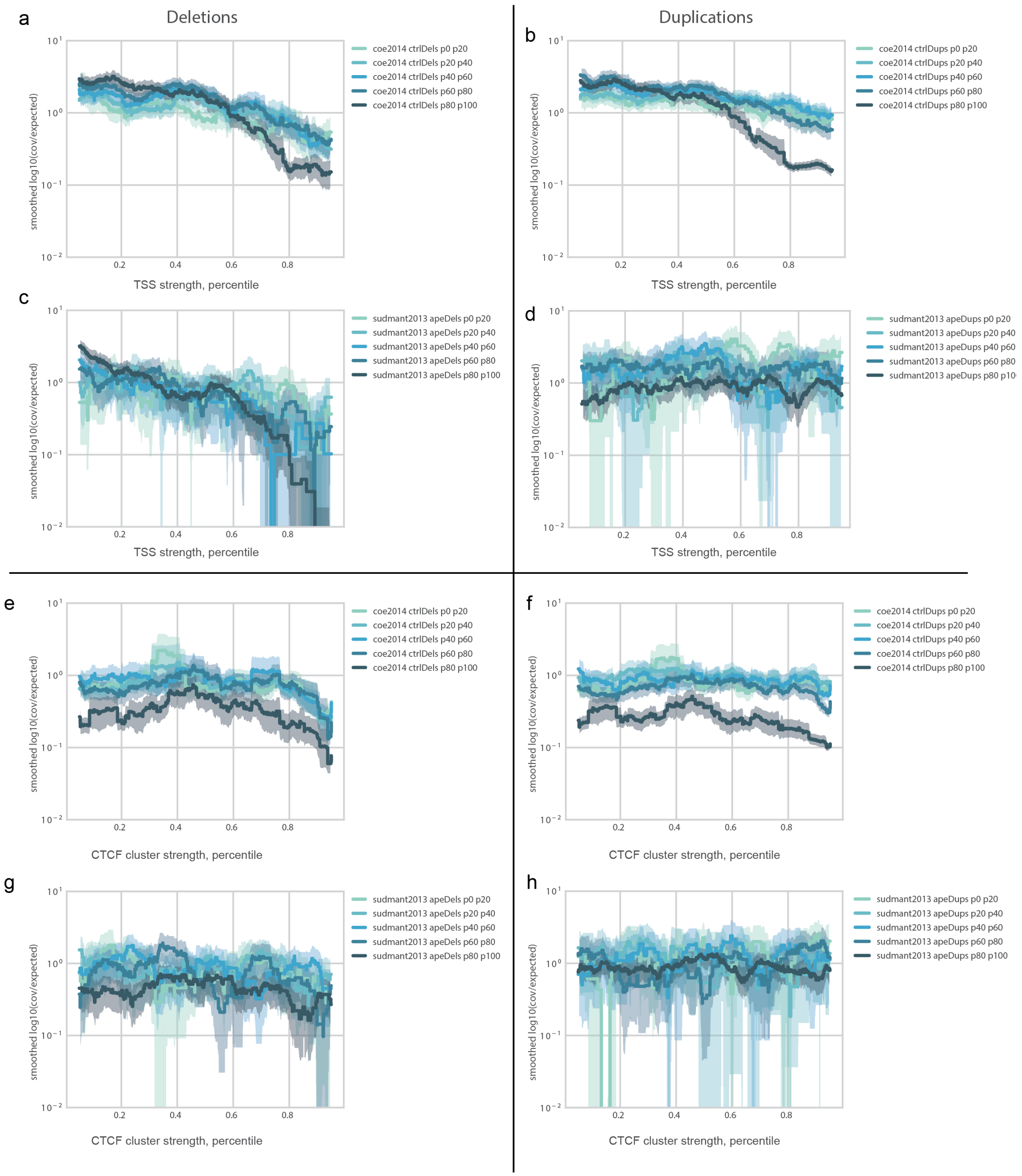


**Supplemental Fig. 4.**

Coverage for deletions (**a,c,e,g**) and duplications (**b, d, f, h**), versus total expression (**a-d**) or CTCF strength (**e-h**), additionally stratified by variant length (five quantiles), for the indicated datasets. Sliding window and bootstraps as in **Supplemental Fig. 3**. Note that the longest duplications are the main contributor to the avoidance of active TSSs and strongly bound CTCF sites observed for healthy human variants.

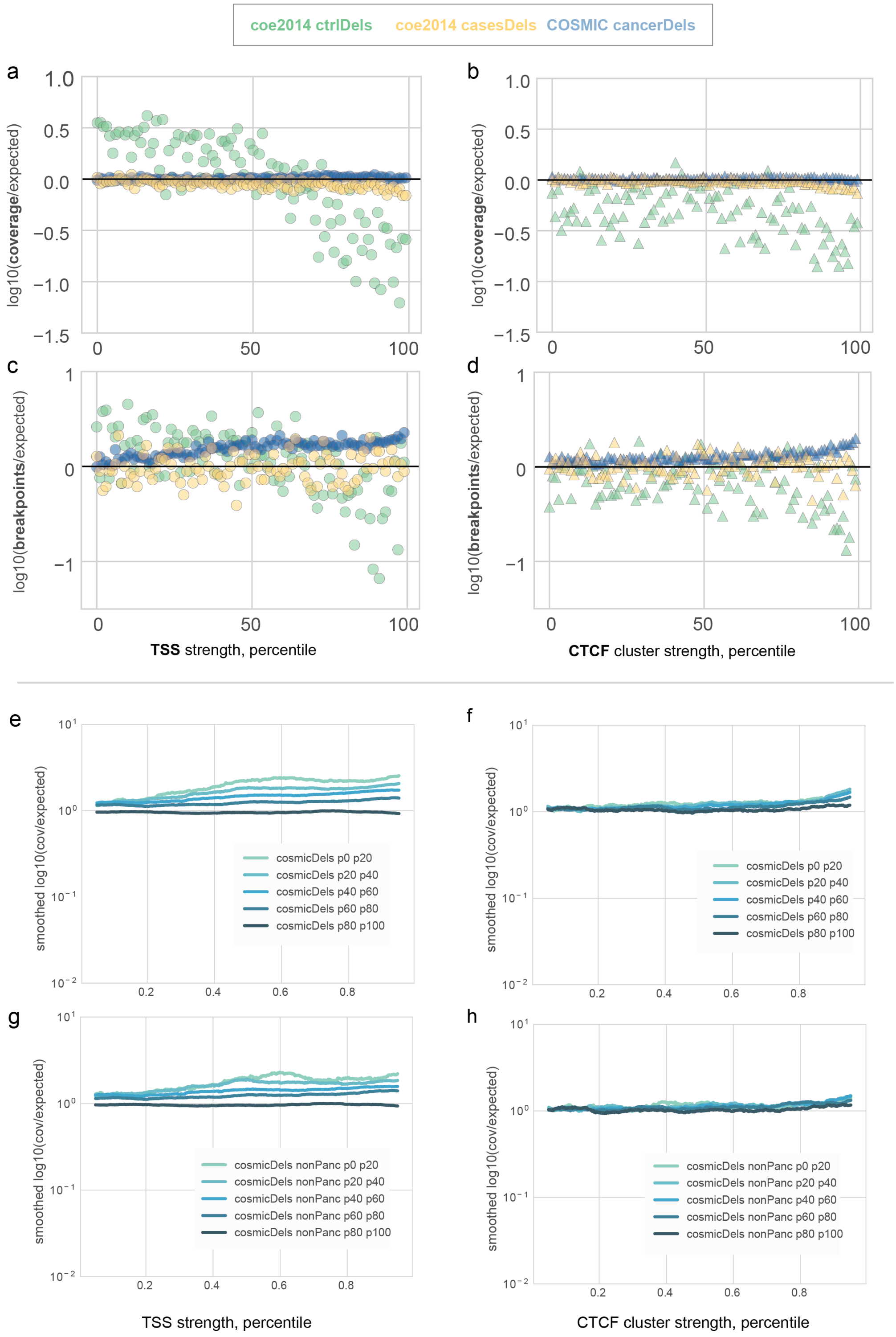


**Supplemental Fig. 5. (caption on next page)**

**Supplemental Fig. 5.**

**a.** Deletion coverage is relatively flat as a function of TSSs strength when considering all deletions across all cancer types in COSMIC.

**b.** Deletion coverage is also relatively flat as a function of CTCF cluster strength

**c.** Deletion breakpoint frequency, however, is highest for the most active TSSs for COSMIC deletions (blue), opposite the trend in healthy humans (green), and distinct from the apparent lack of a trend for deletions in patients with developmental delay and autism (yellow).

**d.** Deletion breakpoint frequency shows a similar trend for CTCF cluster strength.

**e.** Deletion coverage, stratified by deletion length shows short deletions have a greater dependence on TSSs strength, as expected, and explaining the difference between coverage and breakpoint frequency in **a-d**.

**f.** A similar pattern is see as a function of CTCF cluster strength.

**g,h.** As pancreatic cancers contributed roughly half of the deletions in COSMIC, we re-examined the above two trends excluding pancreatic deletions, and saw similar patterns for TSSs and CTCF clusters (as in **e,f**).

**Supplemental Table 1.** GO-rilla enrichments for TSSs around TAD boundaries that were significantly deleted in the ape, healthy human, and patients with developmental disease or autism datasets. All three sets of terms were calculated relative to the set of all TSSs +/-500kb around TAD boundaries using the ‘two unranked lists of genes’ running mode (http://cbl-gorilla.cs.technion.ac.il/).

**Supplemental Table 2.** Input lists for GO-rilla enrichment calculations (three target sets and the background set).
